# PP2A catalytic subunit alpha is critically required for CD8^+^ T cell homeostasis and anti-bacterial responses

**DOI:** 10.1101/2024.02.06.578745

**Authors:** Xian Zhou, Meilu Li, Minji Ai, Yanfeng Li, Xingxing Zhu, Michael J. Hansen, Jun Zhong, Kenneth L. Johnson, Roman Zenka, Akhilesh Pandey, Larry R. Pease, Hu Zeng

**Affiliations:** Division of Rheumatology, Department of Medicine, Mayo Clinic, Rochester, MN 55905, USA; Department of Immunology, Mayo Clinic, 200 First Street SW, Rochester, MN 55905, USA; Department of Laboratory Medicine and Pathology, Mayo Clinic, Rochester, MN 55905, USA; Proteomics Core, Mayo Clinic, Rochester, MN 55905, USA

## Abstract

While the functions of tyrosine phosphatases in T cell biology have been extensively studied, our knowledge on the contribution of serine/threonine phosphatases in T cells remains poor. Protein phosphatase 2A (PP2A) is one of the most abundantly expressed serine/threonine phosphatases. It is important in thymocyte development and CD4^+^ T cell differentiation. Utilizing a genetic model in which its catalytic subunit alpha isoform (PP2A Cα) is deleted in T cells, we investigated its contribution to CD8^+^ T cell homeostasis and effector functions. Our results demonstrate that T cell intrinsic PP2A Cα is critically required for CD8^+^ T cell homeostasis in secondary lymphoid organs and intestinal mucosal site. Importantly, PP2A Cα deficient CD8^+^ T cells exhibit reduced proliferation and survival. CD8^+^ T cell anti-bacterial response is strictly dependent on PP2A Cα. Expression of Bcl2 transgene rescues CD8^+^ T cell homeostasis in spleens, but not in intestinal mucosal site, nor does it restore the defective anti-bacterial responses. Finally, proteomics and phosphoproteomics analyses reveal potential targets dependent on PP2A Cα, including mTORC1 and AKT. Thus, PP2A Cα is a key modulator of CD8^+^ T cell homeostasis and effector functions.

## INTRODUCTION

Protein phosphatase 2A (PP2A) is a ubiquitously expressed serine/threonine phosphatase that controls a plethora of cellular processes. It is a heterotrimeric enzyme composed of a scaffolding subunit known as A or PR65 subunit, a catalytic subunit known as C subunit, and a variable regulatory subunit known as B subunit. While its predicted functions are vast, genetic studies indicate that its functions can be cell type and context specific. In recent years, the importance of PP2A in immunity has been appreciated. Within adaptive immune system, PP2A is essential to support thymocyte survival^1^, promote Th17 differentiation^2,3^, maintain Treg function^4^, and sustain B cell development and activation^5,6^. Yet, overexpression of PP2A catalytic subunit also contributes to development of systemic lupus erythematosus by partly by increasing IL-17 production, suppressing IL-2 production and CD4^+^ follicular helper T cells^7–9^, indicating important functions of PP2A in lymphocyte development, activation and tolerance. Mechanistically, PP2A may function as a suppressor of mechanistic target of rapamycin (mTOR)^4,10,11^. Recent studies proposed that PP2A inhibition could synergize with immune checkpoint inhibition therapy to improve anti-tumor immune response through reduction of regulatory T cells and increased Th1 differentiation^12^. However, its function in mature CD8^+^ T cells remains less understood. While one study indicates that one B subunit, B55b, promotes CD8^+^ T cell apoptosis^13^, the function of C subunit in CD8^+^ T cells remains largely unknown. Moreover, PP2A catalytic subunit alpha (PP2A Cα) appears to protect thymocytes from apoptosis^1^, which contradicts the pro-apoptotic role of B55b^13^.

Here we investigated T cell homeostasis and CD8^+^ T cell effector response in a mouse model with T cell specific deletion of PP2A Cα. Our data demonstrate that PP2A Cα is critically required for mature T cell homeostasis, particularly conventional CD8αβ T cells in the intestine. Contrary to the current model in which PP2A exerts a negative role in T cell activation^14,15^, PP2A Cα deficient CD8^+^ T cells had significantly reduced proliferation and survival. While overexpression of anti-apoptosis molecule Bcl2 rescued peripheral T cell homeostatic defects in PP2A Cα deficient mice, it did not restore CD8αβ T cells in the intestine, suggesting that cell death plays a role in T cell homeostasis in peripheral lymphoid tissue, but not in mucosal site. Furthermore, we showed that PP2A Cα deficient CD8^+^ T cells failed to respond to listeria infection, which is also independent of cell apoptosis. Finally, proteomics and phosphor-proteomics analysis revealed potential signaling events that are dependent on PP2A Cα. Together, our data reveal essential functions of PP2A for CD8^+^ T cell homeostasis and effector activity.

## RESULTS

### PP2A Cα is critically required for T cell homeostasis

To investigate the function of PP2A in mature T cells, we generated *Cd4^Cre^Ppp2ca*^f/f^ mice, in which the Cα subunit was deleted in both CD4^+^ and CD8^+^ T cells starting at CD4/CD8 double positive stage in the thymus. The transcription of *Ppp2ca*, but not *Ppp2cb* (encoding the Cβ subunit), was completely abolished in total splenic T cells (**Fig. 1A**). Consequently, the PP2A catalytic activity was reduced by about 50% in T cells (**Fig. 1B**). Although a previous study using proximal *Lck*-Cre driven deletion of *Ppp2ca* leads to defective thymocyte development, deletion of *Ppp2ca* via *Cd4*-Cre did not have appreciable effects on thymocyte development. The absolute number of thymocytes, and the frequencies of double negative (DN), double positive (DP), CD4^+^ single positive (SP) and CD8^+^ SP cells were equivalent between WT and *Cd4^Cre^Ppp2ca*^f/f^ mice (**Fig. 1C**). In the spleen and lymph nodes, we observed reduced T cell frequencies and total T cell numbers (**Fig. 1D-1F**), indicating that secondary lymphoid tissue T cell homeostasis is compromised in the absence of PP2A Cα. We further examined the naïve and memory T cell subsets by staining CD62L and CD44. The data indicate that while PP2a Cα deficient CD4^+^ T cells had normal distribution of naïve and memory CD4^+^ T cells, *Ppp2ca* deficiency led to a modest reduction of naïve CD8 T cells, and an increase of CD44^hi^CD62L^hi^ central memory phenotype CD8^+^ T cells, compared to control mice (**Fig. 1G**). T cells in mucosal sites, such as small and large intestines, provide barrier defense and have unique phenotypes^16^. In terms of intestinal CD4^+^ T cells, we also observed a significant reduction of CD4^+^ T cells in lamina propria of *Cd4^Cre^Ppp2ca*^f/f^ mice (**Fig. 1H**). Strikingly, we found that the frequency and number of intraepithelial CD8αβ T cells in *Cd4^Cre^Ppp2ca*^f/f^ mice reduced about ten folds compared to that in WT mice, while CD8αα T cells had a modest reduction (two folds or less) in the absence of PP2A Cα (**Fig. 1I**). Finally, to test if the observed T cell homeostasis phenotypes were cell intrinsic, we generated bone marrow (BM) chimeras by mixing BM cells from WT or *Cd4^Cre^Ppp2ca*^f/f^ mice with BM cells from congenically marked CD45.1^+^ mice at 1:1 ratio. Analyses of T cell homeostasis in secondary lymphoid organs (**Fig. 1J**) and intestinal sites (**Fig. 1K**) showed that *Cd4^Cre^Ppp2ca*^f/f^ chimera mice had substantial reduction of CD4^+^ and CD8^+^ T cell frequencies. Altogether, these data demonstrated that PP2A Cα is dispensable for thymocyte development, but it is critically required for mature T cell homeostasis, particularly for intestinal CD8αβ T cells in *Cd4^Cre^Ppp2ca*^f/f^ mice.

**Figure 1.**
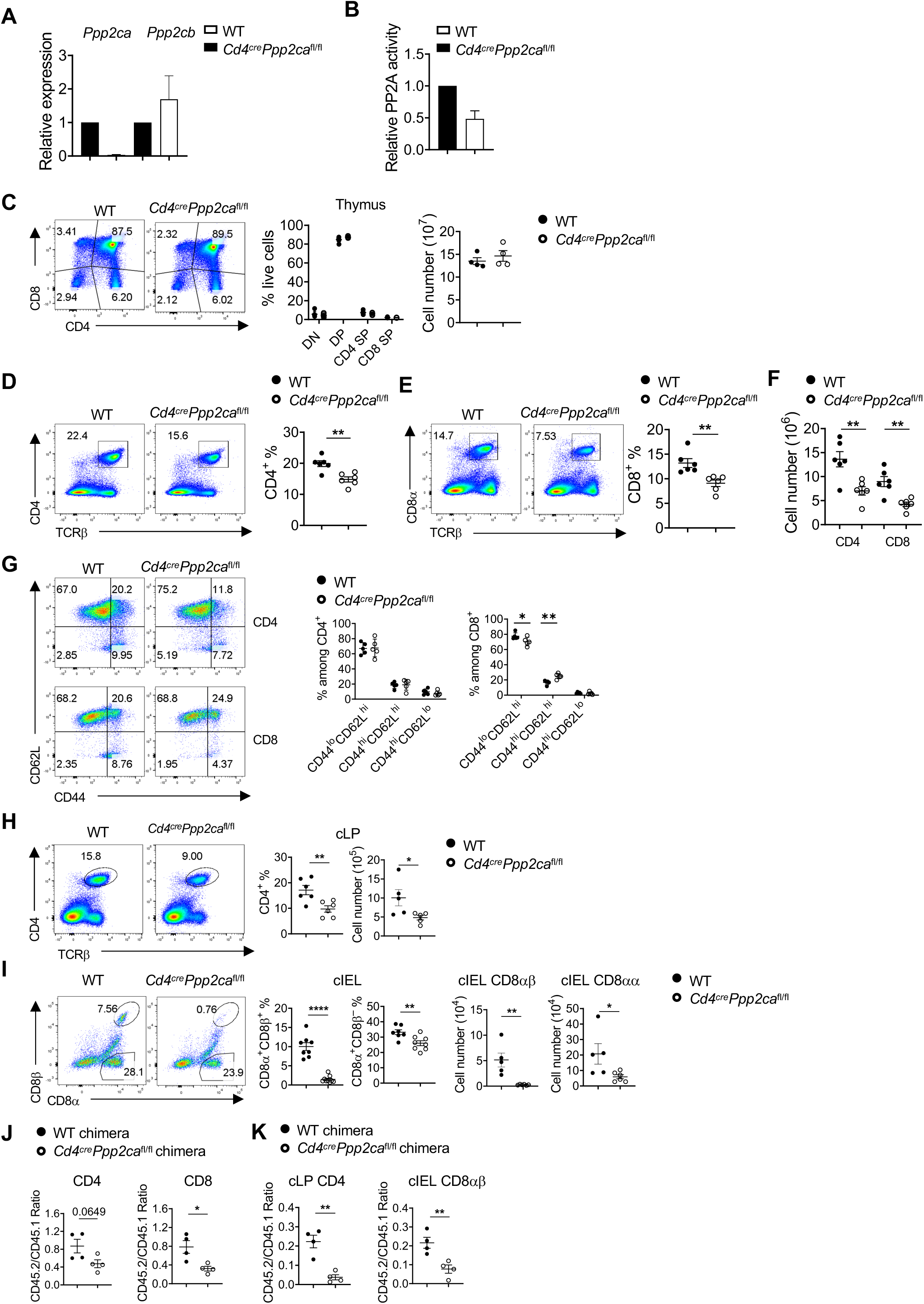
PP2A catalytic subunit alpha is required to maintain peripheral CD4 and CD8 T cells homeostasis. (A) mRNA expression of two isoforms of PP2A catalytic unit, *Ppp2ca* and *Ppp2cb* among total T cells isolated from WT and *Cd4^Cre^Ppp2ca*^fl/fl^ spleens. (B) Phosphatase PP2A activity among total splenic T cells in WT and *Cd4^Cre^Ppp2ca*^fl/fl^. (C) Flow cytometry analysis of thymic T cells, Right, the frequencies and numbers of CD4^−^CD8^−^ double negative (DN), CD4^+^CD8^+^ double positive (DP), CD4 single positive (CD4 SP) and CD8 single positive (CD8 SP) cells in mouse thymus. (D-F) Flow cytometry analysis of splenic T cells. Right, the frequencies of both CD4^+^ (D) and CD8^+^ T cells (E) in mouse spleens. (F) Summary of the numbers of splenic CD4^+^ and CD8^+^ T cells. (G) Expression of CD44 and CD62L in splenic CD4^+^ and CD8^+^ T cells. Right, summary of CD44^lo^CD62L^hi^, CD44^hi^CD62L^hi^ and CD44^hi^CD62L^lo^ percentages among splenic CD4^+^ and CD8^+^ T cells. (H-I) Flow cytometry analysis of gut T cells. (H) The frequency and number of CD4^+^ T cells in mouse colon lamina propria (cLP). (I) The frequencies and numbers of intraepithelial lymphocyte (IEL) conventional CD8 (CD8αβ) and CD8αα T cells in mouse colons. (J-K) T cell maintenance in mixed bone marrow chimera mouse model at homeostasis state. Mixed bone marrow cells (CD45.2^+^) from WT and *Cd4^Cre^Ppp2ca*^fl/fl^ mice were transferred to synthetic irradiated CD45.1^+^ WT recipient mice. Mice were analyzed after bone marrow reconstitution for 8 weeks. (J) The ratio of CD45.2/CD45.1 of both splenic CD4^+^ and CD8^+^ T cells. (K) The ratio of CD45.2/CD45.1 of cLP CD4^+^ T cells and cIEL CD8αβ T cells from mixed bone marrow chimeras. * *P* < 0.05, ** *P* < 0.01, **** *P* < 0.0001. *p* values were calculated with unpaired Student’s *t* test. Results (A-K) were pooled from at least 3 independent experiments. Error bars represent SEM.

### PP2A supports peripheral, but not mucosal, T cell homeostasis partly through enhancing survival

Peripheral T cells undergo homeostatic proliferation supported by cytokines such as IL-7 and IL-15. We hypothesized that the substantial reduction of T cell homeostasis in the absence of PP2A Cα might be due to impaired homeostatic proliferation. To test this, we mixed T cells from WT or *Cd4^Cre^Ppp2ca*^f/f^ mice with CD45.1 congenic T cells at 1:1 ratio, and transferred them into *Rag1^−/−^* mice. WT T cells underwent vigorous homeostatic proliferation, but PP2A Cα deficient T cells showed reduced proliferation and the ratio between PP2A Cα deficient T cells and CD45.1 T cells reduced to less than 1:10, indicating that PP2A Cα deficiency profoundly impairs the T cell homeostatic proliferation (**Fig. 2A, B**). In addition to reduced homeostatic proliferation, PP2A Cα deficient T cells had a reduced proliferation in a mixed lymphocyte reaction system (**Fig. 2C**). When stimulated with plate coated anti-CD3/anti-CD28, PP2A Cα deficient CD8^+^ T cells showed a relatively normal cell division, but consistently reduced cell recovery (**Fig. 2D**), suggesting an increased cell death. To further examine apoptosis, we performed Annexin-V staining *in vitro* stimulated T cells. We observed a modest increase of apoptosis in PP2A Cα deficient CD8^+^ T cells (**Fig. 2E**). Moreover, we also found that PP2A Cα deficient T cells had increased level of cleaved-caspase 3, a direct indication of apoptosis (**Fig. 2F**). These observations are consistent with a recent study indicating that PP2A is required to prevent thymocyte apoptosis^1^. Thus, PP2A Cα is required for T cell proliferation and survival *in vitro* and *in vivo*.

**Figure 2.**
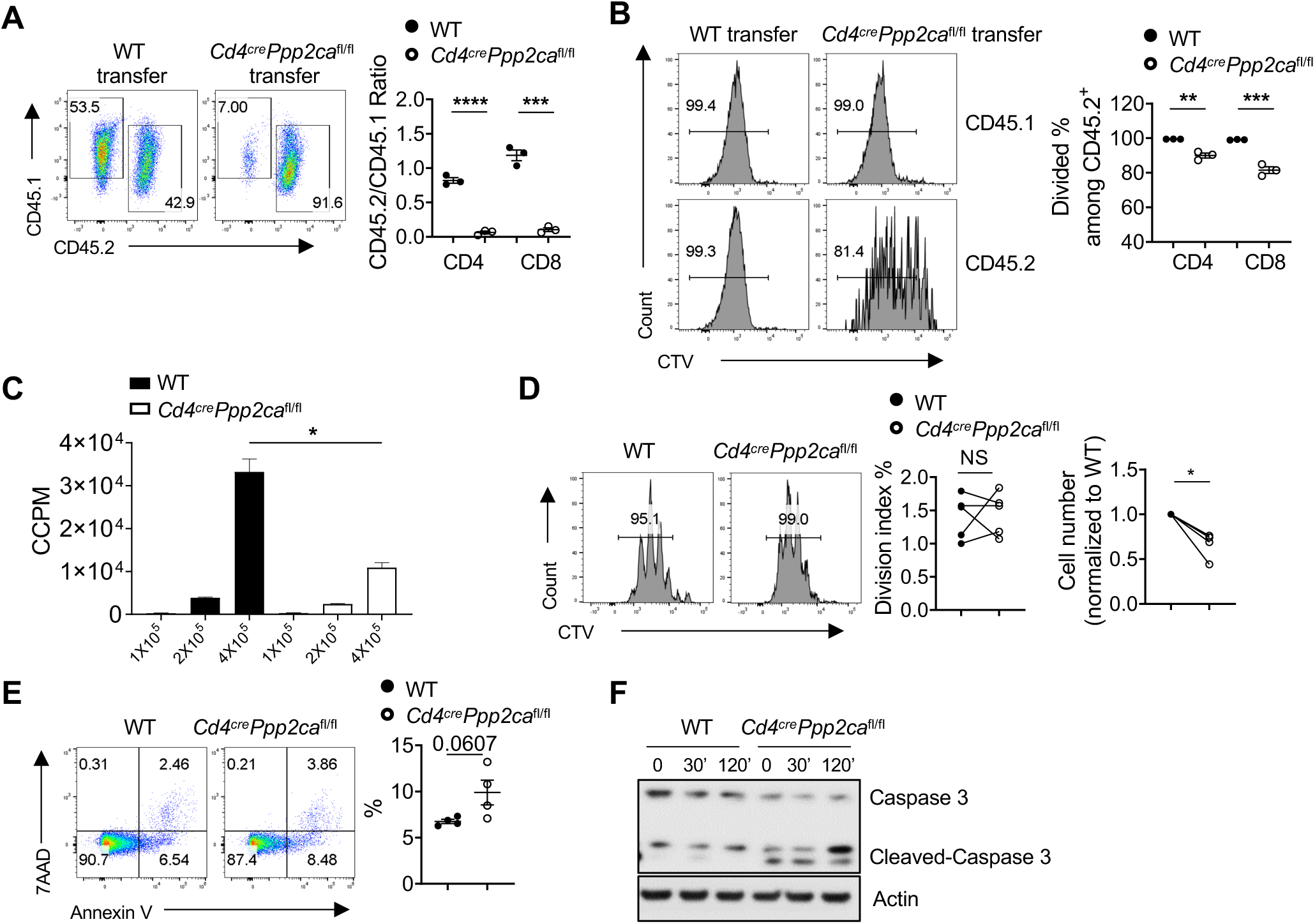
Loss of PP2A Cα reduces T cell proliferation and survival *in vivo* and *in vitro*. (A-B) Total splenic T cells (CD45.2^+^) were purified from WT and *Cd4^Cre^Ppp2ca*^fl/fl^ mice, 1:1 mixed with purified CD45.1^+^ WT T cells, labeled with CellTrace Violet (CTV) dye, and transferred into *Rag1^−/−^*mice. One week after transfer, flow analysis of the T cell maintenance and division *in vivo*. (A) Flow analysis of T cell maintenance. Right, the ratio of CD45.2/CD45.1 of both splenic CD4^+^ and CD8^+^ T cells. (B) Flow analysis of T cell division after transfer into *Rag1^−/−^*mice. Right, the percentages of divided splenic CD4^+^ and CD8^+^ T cells. (C) T cells isolated from WT and *Cd4^Cre^Ppp2ca*^fl/fl^, at indicated numbers, were stimulated with irradiated splenocytes from allogenic Balb/c mice. After culturing for 3 days, the mixed lymphocyte reaction (MLR) was measured with [^3^H]-thymidine incorporation assay. (D) Naïve CD8^+^ T cells purified from WT and *Cd4^Cre^Ppp2ca*^fl/fl^ were labeled with CTV and stimulated with anti-CD3/anti-CD28 plus IL-2. Right, The division and live cell number of T cells after 3-days culture. (E) Analysis of early apoptotic (Annexin V^+^7AAD^−^) T cells percentage isolated from WT and *Cd4^Cre^Ppp2ca*^fl/fl^ stimulated with anti-CD3/anti-CD28 plus IL-2 overnight. (F) CD8^+^ T cells from WT and *Cd4^Cre^Ppp2ca*^fl/fl^ mice were stimulated with anti-CD3/anti-CD28 plus IL-2 for 3 days, rested and restimulated with IL-2. Expression of caspase-3 and cleaved caspase-3 was examined by immunoblot. NS, not significant; * *P* < 0.05, ** *P* < 0.01, *** *P* < 0.001, **** *P* < 0.0001. *p* values were calculated with unpaired Student’s *t* test. Results were presentative of two (A-B) and three (C, F), or were pooled from at least 3 (D-E) independent experiments. Error bars represent SEM.

To further investigate whether suppression of apoptosis may rectify the defects found in *Cd4^Cre^Ppp2ca*^f/f^ mice, we bred *Cd4^Cre^Ppp2ca*^f/f^ mice with Bcl2 transgenic (Bcl2-Tg) mice, which could rectify increased cell death in T cells^17^. Analysis of the T cell homeostasis in WT, *Cd4^Cre^Ppp2ca*^f/f^, Bcl2-Tg, and *Cd4^Cre^Ppp2ca*^f/f^; Bcl2-Tg mice revealed that Bcl2 transgene successfully rescued the CD4^+^ and CD8^+^ T cell frequencies in secondary lymphoid tissues (**Fig. 3A**). However, Bcl2 transgene did not restore T cell homeostasis in intestinal lamina propria and among intraepithelial lymphocytes (**Fig. 3B, 3C**). These data indicate that increased apoptosis could underly the reduced T cell homeostasis in secondary lymphoid tissues, but likely not in intestinal sites, in *Cd4^Cre^Ppp2ca*^f/f^ mice.

**Figure 3.**
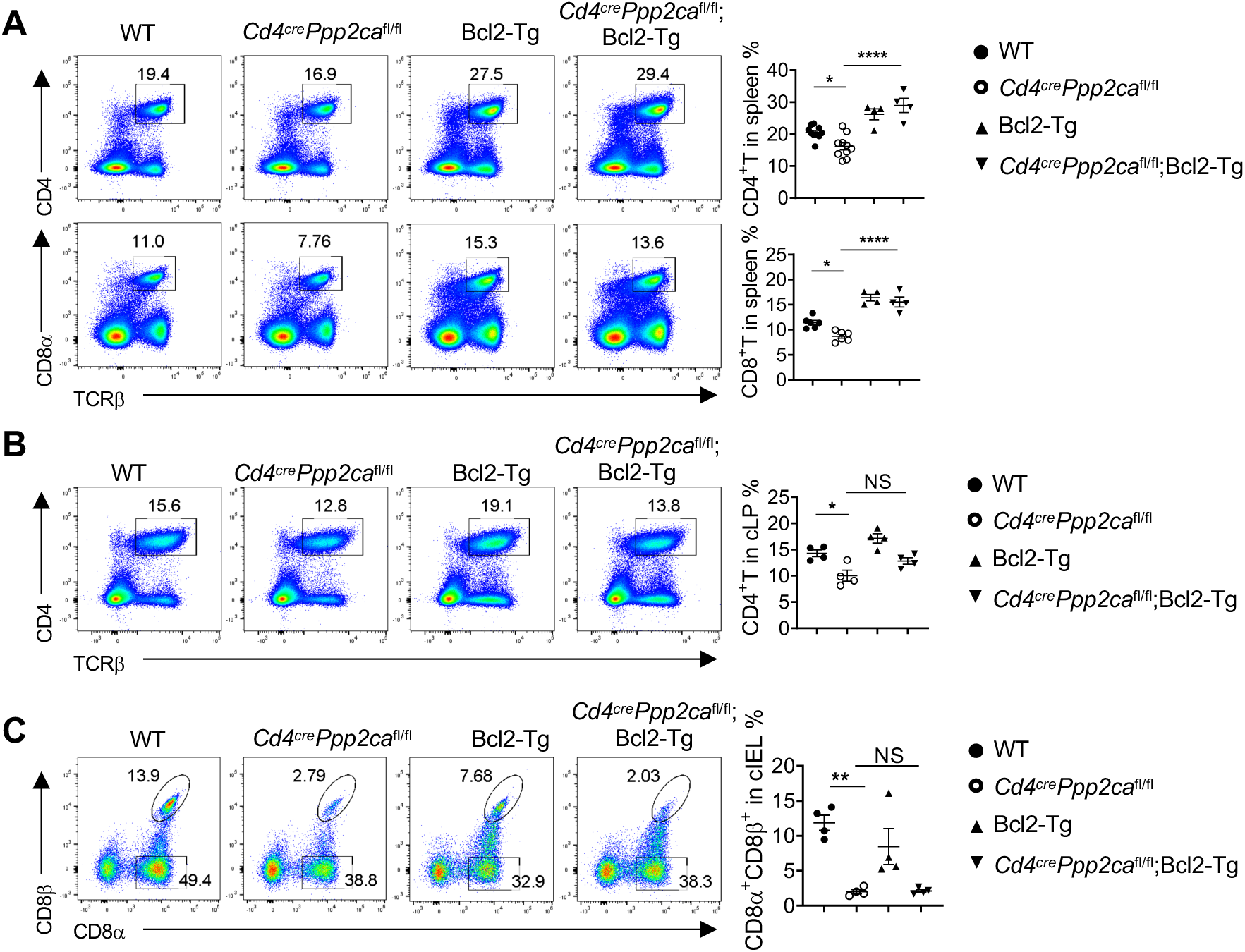
Overexpression of Bcl2 can rescue the loss of splenic T cells, but not gut T cell homeostasis after *Ppp2ca* deletion. (A) Flow analysis of both splenic CD4^+^ and CD8^+^ T cells from WT, *Cd4^Cre^Ppp2ca*^fl/fl^, Bcl2*-*Tg and *Cd4^Cre^Ppp2ca*^fl/fl^; Bcl2-Tg mice. Right, summary of the percentages of splenic CD4^+^ (upper) and CD8^+^ (lower) T cells. (B-C) Flow analysis of colon lamina propria (cLP) CD4^+^ and CD8^+^ T cells from WT, *Cd4^Cre^Ppp2ca*^fl/fl^, Bcl2*-*Tg and *Cd4^Cre^Ppp2ca*^fl/fl^; Bcl2-Tg mice. Right, the summary of percentages of cLP CD4^+^ T cells (B) and colon intraepithelial conventional CD8αβ T cells (C) at homeostasis state. NS, not significant; * *P* < 0.05, ** *P* < 0.01, **** *P* < 0.0001 (one-way ANOVA). Results were pooled from at least 3 independent experiments. Error bars represent SEM.

### CD8^+^ T cell anti-bacterial effector function depends on PP2A Cα

We sought to further evaluate the *in vivo* impact of *Ppp2ca* deficiency on T cell functions. To this end, we infected WT and *Cd4^Cre^Ppp2ca*^f/f^ mice with *Listeria monocytogene* expressing OVA (Lm-OVA). One week after infection, we found that there was a substantial reduction of CD8^+^ T cell frequencies in spleen and liver (**Fig. 4A**). Staining with OVA tetramer showed drastically reduced antigen specific CD8^+^ T cells in spleen and liver (**Fig. 4B**). Furthermore, PP2A Cα deficient CD8^+^ T cells produced significantly reduced IFNγ and TNFα upon stimulation with OVA_257-264_ peptide, indicating reduced antigen specific effector activity (**Fig. 4C**). To test if the reduced effector T cell generation was intrinsic to lymphocytes, we infected WT and *Cd4^Cre^Ppp2ca*^f/f^ chimera mice with Lm-OVA. *Ppp2ca* deficient CD8^+^ T cells expanded less efficiently (**Fig. 4D**), generated significantly reduced OVA specific CD8^+^ T cells (**Fig. 4E**) and OVA specific IFNγ and TNFα production (**Fig. 4F**), demonstrating that these phenotypes were immune cell intrinsic. Finally, we found that Bcl2-Tg failed to restore the defective CD8^+^ T cell responses to Lm-OVA infection in the absence of PP2A Cα (**Fig. 4G and 4H**). Thus, PP2A Cα deficient T cells failed to generate antigen specific CD8^+^ T cell response upon listeria infection independent of their survival defects.

**Figure 4.**
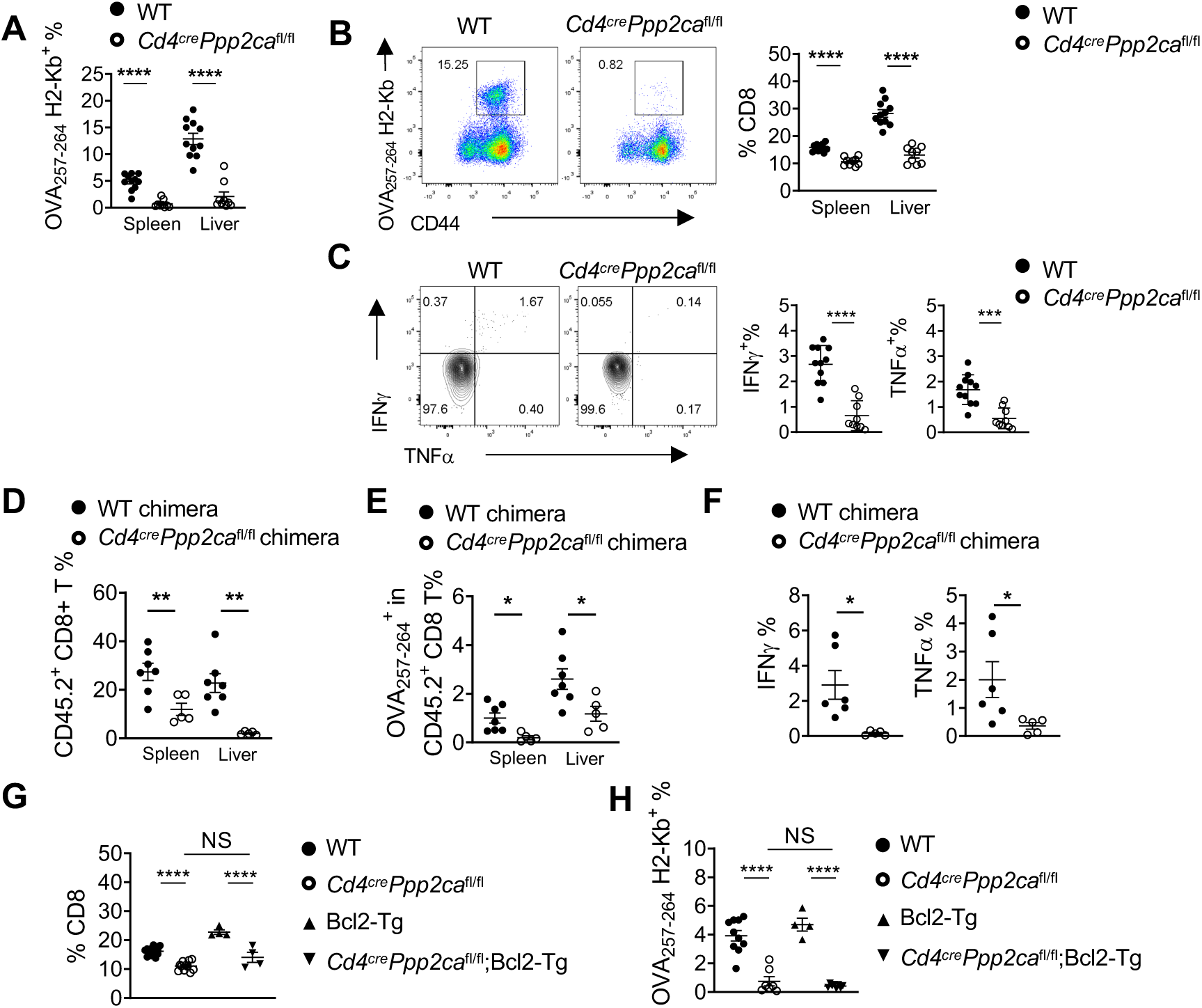
PP2A Cα controls antigen specific T cell response during *Listeria monocytogenes* infection. (A) Summary of the frequencies of total CD8^+^ T cells in both spleen and liver after *Listeria monocytogenes* (LM) intravenous infection in WT and *Cd4^Cre^Ppp2ca*^fl/fl^ mice for 7 days. (B) Frequencies of OVA_257-264_-specific CD8^+^ T cells after infection. Right, summary of the percentage of antigen specific CD8 T cells in both spleen and liver. (C) Flow cytometry analysis of cytokine expression among total splenic CD8^+^ T cells after OVA_257-264_ peptide stimulation for 5 hours. Right, the percentages of IFNγ^+^ and TNFα^+^ CD8 T cells, respectively. (D-F) LM infection in WT and *Cd4^Cre^Ppp2ca*^fl/fl^ mixed bone marrow chimera mice for 7 days. (D) Summary of the percentages of CD45.2^+^ CD8^+^ T cells among total CD8^+^ T cells in spleen and liver after infection. (E) Summary of the percentages of OVA antigen specific CD8 T cells among whole CD45.2^+^ CD8^+^ T cells. (F) Percentages of IFNγ^+^ and TNFα^+^ among CD45.2^+^ CD8^+^ T cells from WT and *Cd4^Cre^Ppp2ca*^fl/fl^ chimera mice after LM infection. (G) Percentages of CD8^+^ T cells in WT, *Cd4^Cre^Ppp2ca*^fl/fl^, Bcl2-Tg and *Cd4^Cre^Ppp2ca*^fl/fl^; Bcl2-Tg mice in spleen after LM infection. (H) Summary of the percentages of OVA_257-264_-specific CD8^+^ T cells in WT, *Cd4^Cre^Ppp2ca*^fl/fl^, Bcl2*-*Tg and *Cd4^Cre^Ppp2ca*^fl/fl^; Bcl2-Tg mice after LM infection, respectively. NS, not significant; * *P* < 0.05, ** *P* < 0.01, **** *P* < 0.0001. (A-F, unpaired Student’s *t* test, G-H, one-way ANOVA). Results were presentative of three (D-F) and two (G, H), or pooled from three (A-C) independent experiments. Error bars represent SEM.

### Multiple pathways were disturbed in *Ppp2ca* deficient CD8^+^ T cells

To further examine the molecular alterations in the PP2A Cα deficient CD8^+^ T cells, we combined tandem-mass-tag (TMT)-based quantitative analysis with immobilized metal affinity chromatography (IMAC)-based phosphopeptide enrichment to examine the changes of proteomes and phosphoproteomes in anti-CD3 and anti-CD28 stimulated CD8^+^ T cells from *Cd4^Cre^Ppp2ca*^f/f^ mice verses WT mice. As shown in **Figure 5A**, we identified 41 proteins whose expression had more than 50% reduction compared to WT cells, and 44 proteins that had increased more than 50% (p value =< 0.05). As expected, the most downregulated protein was PP2A Cα. Other significantly reduced proteins include other PP2A components, such as PPP2CB (the beta isoform of PP2A catalytic subunit), PPP2R5A, PPP2R5B, PPP2R1A, and PPP2R5C, suggesting that deficiency of PP2A Cα might destabilize PP2A holoenzyme (**Fig. 5A**). When we examined the significantly altered phosphorylation sites, we observed that several serine residues on Rps6 had increased phosphorylation in PP2A Cα deficient T cells (**Fig. 5B**), suggesting activation of mTORC1. On the other hand, serine phosphorylation of AKT and tyrosine phosphorylation of Stat5 were reduced in the absence of PP2A Cα, which could be a secondary effect of PP2A Cα deficiency because PP2A is a Ser/Thr phosphatase. Ingenuity pathway analysis (IPA) using phosphoproteomics data identified PI3K/AKT pathway and EIF2 signaling among the top pathways associated with *Ppp2ca* deficiency (**Fig. 5C**). Immunoblot assay confirmed a modest increase of S6 phosphorylation and reduced AKT phosphorylation in PP2A Cα deficient CD8^+^ T cells (**Fig. 5D**). mTORC1 is a central metabolic regulator during lymphocyte activation and overactivation of mTORC1 leads to the loss of quiescence and cell death in CD8^+^ T cells^17,18^. Consistent with the modest increase of mTORC1 activation and cell apoptosis in the absence of PP2A Cα, we observed increased basal oxygen consumption rate in PP2A Cα deficient CD8^+^ T cells, suggesting that PP2A Cα may control CD8^+^ T cell metabolism by suppressing mTORC1 (**Fig. 5E**). Altogether, these data indicate that PP2A Cα contributes to CD8^+^ T cell functions through multiple pathways.

**Figure 5.**
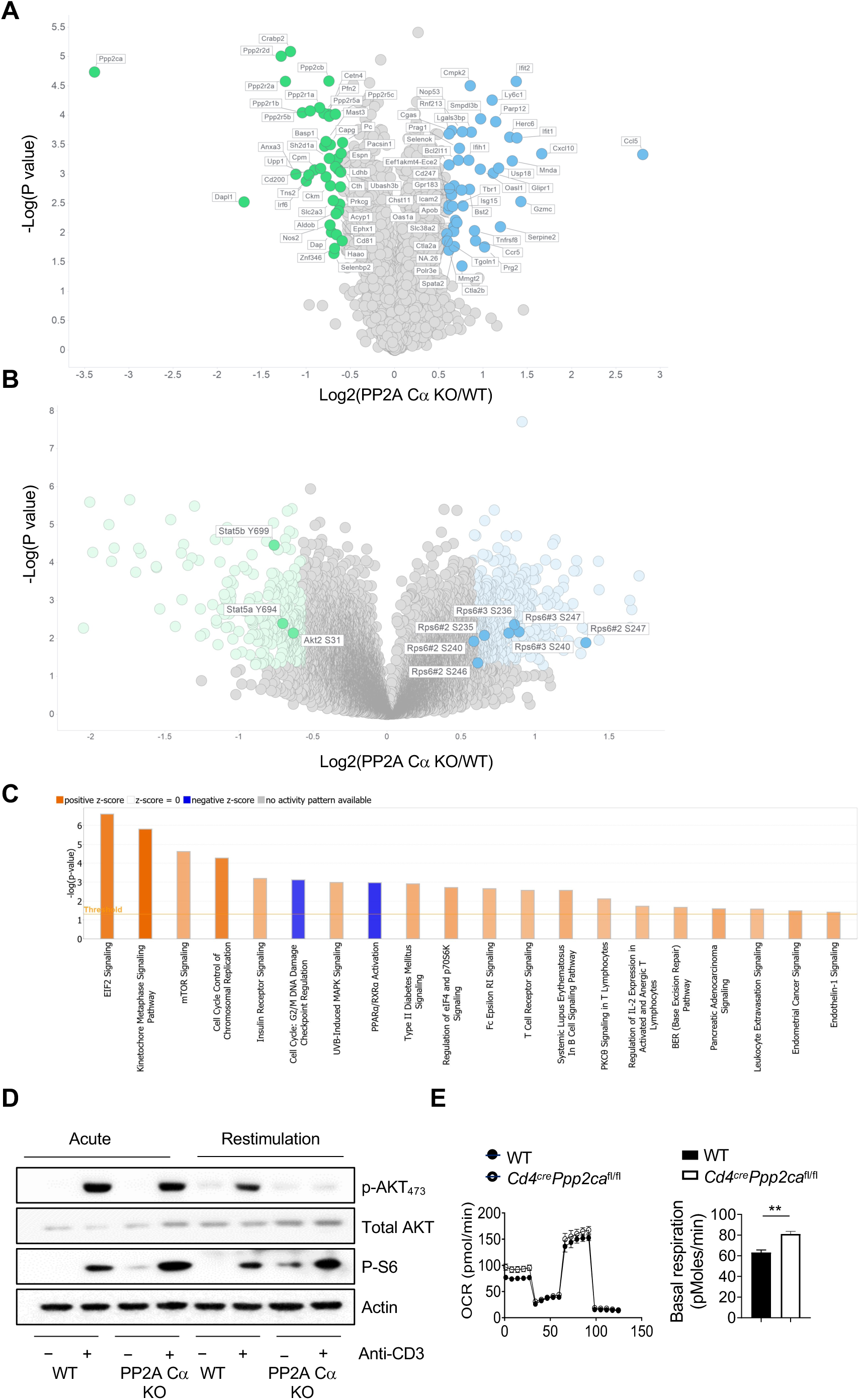
Proteomics analysis of WT and PP2A Cα deficient CD8^+^ T cells. Volcano plots depicting of the up-regulated and down-regulated proteins (A) and phosphorylation sites (B) in anti-CD3/anti-CD28 stimulated CD8^+^ T cells from *Cd4^Cre^Ppp2ca*^fl/f^ mice verses WT mice (Log2FC cutoff: > |±0.585|, −Log10 P value cutoff: > 1.3). (C) Ingenuity pathway analysis (IPA) of phosphoproteomics data for CD8^+^ T cells from *Cd4^Cre^Ppp2ca*^fl/fl^ mice verses WT mice. (D) Immunoblot analysis of *in vitro* stimulated CD8^+^ T cells. Briefly, naïve CD8^+^ T cells were isolated from WT and *Cd4^Cre^Ppp2ca*^fl/fl^ mice. For acute stimulation, naive CD8^+^ T cells were stimulated with anti-CD3 crosslink plus anti-CD28 for 5 minutes. For restimulation, naïve CD8^+^ T cells were cultured with coated anti-CD3/anti-CD28 plus IL-2 for 3 days, and washed with fresh complete medium and rested for 3 hours, and re-stimulated with crosslink anti-CD3 plus anti-CD28 for 5 minutes. The p-AKT_473_, total AKT, p-S6 were determined by immunoblot. (E) Naïve CD8^+^ T cells were activated with coated anti-CD3/anti-CD28 plus IL-2 overnight. Oxygen consumption rate (OCR) was measured on a Seahorse XFe96 bioanalyzer using Mitostress assay kit. Right, summary of the basal OCR. ** *P* < 0.01 (unpaired Student’s *t* test). Results were presentative of three (D, E) independent experiments. Error bars represent SEM.

## DISCUSSION

The current model of T cell signaling involves a well-orchestrated network of kinases and phosphatases. Majority of these kinase and phosphatases target tyrosine, such as Src family kinases and Syk family kinases, or lipid, such as PI3K. The functions of serine-threonine kinases have been much less understood or studied. Earlier studies identified PP2A as a binding partner for CTLA-4, but how it regulates CTLA-4 activity remains unclear^14,19^. Previous literatures have suggested that PP2A may suppress T cell anti-tumor effector function and targeting PP2A has been proposed to enhance tumor immunotherapy^10,12,20^. However, PP2A is also a well-known tumor suppressor, which could complicate such approach^21^. Recent genetic mouse studies unveil key functions of PP2A in T cell development and CD4^+^ T cell differentiation^1,2,9^. Here, our genetic study investigated the *in vivo* functions of the catalytic subunit alpha of PP2A for CD8^+^ T cell homeostasis and effector functions. The current model posits that PP2A negatively controls T cell activation partly through binding with CTLA-4 and modulates PI3K-Akt activation^14,15^. Thus, one would expect to observe CD8 T cell overactivation in the absence of PP2A Cα. Yet, our data show that PP2A Cα deficient T cells have reduced proliferation and survival *in vitro* and *in vivo*. Importantly, the diminished T cell functions are likely independent of cell survival defect because overexpression of Bcl2 failed to rectify the reduced anti-bacterial effector T cell generation, even though it could restore CD8 T cell homeostasis in spleen. Furthermore, our data is the first to reveal a critical role of PP2A Cα to maintain mucosal CD8^+^ T cell homeostasis. How PP2A may contribute to mucosal immune response warrants further investigation. Finally, our results indicate non redundant functions between the alpha and beta isoforms of catalytic subunit, because PP2A Cα deficient CD8 T cells preserve about 50% of the catalytic activity, presumably from PP2A Cβ. Future studies are needed to understand the contribution of PP2A Cβ and any redundant and nonredundant functions between the 2 isoforms. Altogether, our study provides definitive evidence that PP2A Cα is critically required for CD8^+^ T cell mucosal homeostasis and anti-bacterial effector functions.

## MATERIALS and METHODS

### Mice

*Cd4^Cre^Ppp2ca^fl/fl^* and BCL2-Tg mice have been described previously^17,22^. CD45.1 and CD45.2 C57BL/6, Balb/c, and *Rag1*^−/−^ mice were purchased from the Jackson Laboratory. Mice were bred and maintained under specific pathogen-free conditions in the Department of Comparative Medicine of Mayo Clinic Rochester. All animal procedures were approved by Institutional Animal Care and Use Committee of Mayo Clinic Rochester.

### T cell isolation

T cell isolation for real-time PCR, immunoblot analysis, *in vitro* culture, and proteomics analysis experiments was performed using magnetic separation from Life Technologies. Naive CD8^+^ T cell isolation was performed with magnetic separation (EasySep™ Mouse Naïve CD8^+^ T Cell Isolation Kit, StemCell Technologies).

### *In vitro* T cell culture

Naive CD8^+^ T cells were stained with CellTrace Violet proliferation dye (Life Technologies) in the presence of soluble IL-2 and plate-bound anti-CD3 (5 μg/ml; 145-2C11; BioXcell)/anti-CD28 (2 μg/ml; 37.51; BioXcell). T cell proliferation was assessed by flow cytometry and cell number were counted with 0.4% trypan blue at the end of the 3-day culture.

### PP2A activity determination

PP2A enzymatic activity was assessed using the PP2A Immunoprecipitation Phosphatase Assay Kit (17-313; Millipore) per the manufacturer’s instructions. Activated T cells were collected after culture for 3 days. Protein extracts were immunoprecipitated with a PP2A C-specific antibody (1D6; Millipore). An appropriate phosphorylated peptide (amino acid sequence K-R-pT-I-R-R, where ‘pT’ indicates phosphorylated threonine) was added to the immunoprecipitated immunocomplexes as a substrate for PP2A C, and samples were incubated at 4 °C in a shaking incubator overnight. Supernatants were transferred to a 96-well plate and released phosphate was measured by the addition of 100 μl malachite green phosphate detection solution in a 15-min colorimetric reaction. Phosphate concentrations were calculated from a standard curve created using serial dilutions of a standard phosphate solution.

### Flow cytometry

Single-cell suspension was prepared by passing spleen and peripheral lymph nodes through a 70-μm nylon mesh. For analysis of surface markers, cells were stained in PBS containing 1% (w/v) BSA on ice for 30 min, with BV510-labeld anti-CD8α (clone:53-6.7; BioLegend), Super Bright 600-labeled anti-CD4 (Clone: SK-3; eBioscience), APC-Cy7-labeled TCRβ (clone: H57-597; BioLegend), APC-labeled anti-CD127 (A7R34; Biolegend), PE-Cy7-labeled anti-KLRG-1 (MAFA; Biolegend), BV711-labeled anti-CD44 (IM7; Biolegend), PE-eflour610-labeled anti-CD62L (MEL-14; eBioscience), Alexa fluor 700-labeled anti-CD8β (Lyt-3; Biolegend), BV650-labeled anti-CD45.1 (Ly5.1; Biolegend), PE-cy7-labeled anti-CD45.2 (Ly5.2; Biolegend). PE labeled OVA_257-264_ H2-Kb tetramer is a gift from NIH tetramer core. Cell viability was assessed using the Fixable Dye Ghost 540 (Tonbo Bioscience), or 7-AAD. Cell apoptosis was determined with FITC-Annexin V/7-AAD kit followed by the manufacture’s protocol (eBioscience).

Splenocytes and peripheral lymph node cells were cultured for 5 h at 37°C in complete medium (1640 RPMI + 10% FBS) plus 0.5 μM OVA_257-264_ and Monensin solution (1,000×; Biolegend) for cytokines detection. Intracellular cytokine staining of APC-labeled anti-IFNγ (clone: XMG1.2; eBioscience) and FITC-labeled anti-TNFα (clone: MP6-XT22; BioLegend) were performed using the BD Cytofix/Cytoperm. Flow cytometry was performed on the Attune NxT (ThermoFisher) cytometer, and analysis was performed using FlowJo software (Tree Star).

### *Rag1*^−/−^ mice transfer

Transfer models were generated by transferring purified CD45.2^+^ WT or *Cd4^Cre^Ppp2ca^fl/fl^* T cells, mixing with 2 million CD45.1^+^ T cells at 1:1 ratio and labeling with CellTrace Violet dye (Invitrogen), into *Rag1*^−/−^ mice. One week after transfer, the mice were sacrificed, and the spleens were examined.

### Mixed lymphocyte reaction

Splenic T cells were isolated from either WT or *Cd4^Cre^Ppp2ca^fl/fl^*mice as responder cells. The spleen cells isolated from allogenic mouse Balb/c were irradiated and used as stimulator cells. The mixed lymphocyte reaction was performed by seeding 1, 2, or 4×10^5^ responder cells into the round-bottom 96-well plate with 4×10^5^ stimulator cells for 3 days. Each group set three repetitions. The reaction was measured by [^3^H]-thymidine incorporation (1 μCi/mL) (American Radiolabeled Chemicals) after pulsing for additional 18 hours.

### Immunoblotting

For immunoblotting, cells were lysed in lysis buffer with protease and phosphatase inhibitors (Sigma-Aldrich). Protein concentration in samples was quantified by BCA assay (Thermo Fisher Scientific) before loading the samples for electrophoresis and membrane transfer. The transferred membrane was blocked with TBST (0.1% Tween 20) containing 5% BSA for 1 h at room temperature. The membrane was incubated with primary antibodies overnight including anti-Cleaved Caspase 3 (9662S; Cell Signaling), anti-p-AKT (Ser473, D9E; Cell Signaling), anti-AKT (pan, C67E7; Cell Signaling), anti-p-S6 (Ser235/Ser236, D57.2.2E; Cell Signaling), anti-PP2A Cα/μ (F-8; Santa Cruz) and anti-β-actin (clone: 13E5; Sigma-Aldrich). Then, the membrane was washed and incubated with the corresponding secondary antibody for subsequent enhanced chemiluminescence (ECL; Thermo Fisher) exposure. The band intensity of all the immunoblot was analyzed by ImageJ software.

### Quantitative Real-time PCR

For mRNA analysis, total mRNA was isolated from mouse T cells by RNeasy Micro kit (Qiagen), reverse transcribed from mRNA to cDNA for subsequent real-time PCR analysis. *Ppp2ca* and *Ppp2cb* in mouse T cells were measured by real-time PCR with a Bio-Rad Realtime PCR system. Beta-actin expression was used as control. The primers information was provided in the following table.

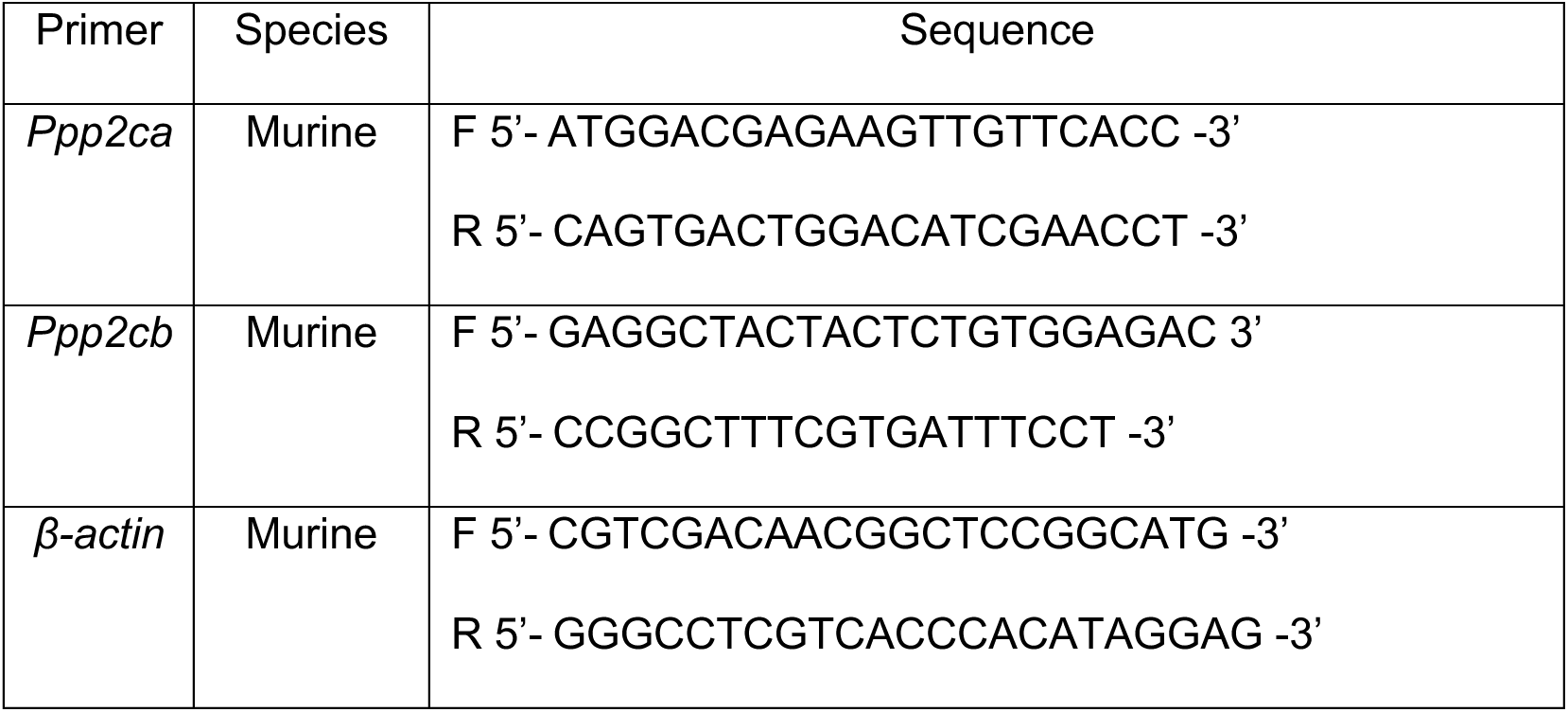

### Mixed bone marrow chimera generation

Chimera mice were constructed by transferring bone marrow cells from WT or *Cd4^Cre^Ppp2ca^fl/fl^*mice, together with bone marrow cells from CD45.1 mice, into lethally irradiated CD45.1 recipients. After 8 weeks of reconstitution, chimera mice were sacrificed, and the spleens and other organs were examined.

### *Listeria monocytogenes* infection

Listeria monocytogenes (LM) derived from the wild type 10403S background deficient in the actin assembly-inducing protein (ΔactA) and expressing amino acids spanning 138-338 of full-length ovalbumin (L. monocytogenes ΔactA-Ova) was obtained from Dr. Brian Sheridan (Stony Brook University)^23^. Bacteria were cultured in BBL brain heart infusion medium (Becton, Dickinson and Company, Sparks, MD) at 37°C, washed, and re-suspended in 1×PBS. In vivo infections were conducted via intravenous (IV) injection of 3×10^4^ colony-forming units (CFU) L. monocytogenes ΔactA-Ova.

### Cell lysis, trypsin digestion, and phosphopeptide enrichment

CD8^+^ T cells isolated from WT or *Cd4^Cre^Ppp2ca*^f/f^ mice were stimulated with anti-CD3 and anti-CD28 in biological triplicates. Stimulated cells were lysed in lysis buffer (9 M urea, 50 mM triethylammonium bicarbonate (TEABC), 1 mM sodium orthovanadate, 1 mM β-glycerophosphate, 2.5 mM sodium pyrophosphate) by sonication on a Bioruptor (Diagenode, Denville, NJ) with a five-minute method of 30s sonication followed by 30s off, at 15°C. After centrifugation, the protein extracts from each sample were reduced with 5 mM dithiothreitol, alkylated with 10 mM iodoacetamide, and subjected to trypsin digestion overnight at ambient temperature. After desalting and lyophilization, the resulting peptides were reconstituted in 100 mM TEABC buffer, pH 8.5. 300 µg of peptide from each sample were labeled with a TMT-6plex reagent, separately. After the confirmation of the labeling efficiency >95%, the excess TMT labeling reagents were quenched with 5% hydroxylamine. The TMT-labeled peptides were mixed and fractionated by basic pH reversed-phase chromatography (bRPLC) to 96 fractions, which then were concatenated to 12 fractions. 5% of each fraction of peptides were aliquoted for LC-MS/MS analysis for global proteomic analysis. The 95% of each fraction of peptides were enriched for phosphopeptides using Fe-NTA cartridges on an Agilent AssayMap Bravo. The enriched phosphopeptides were eluted with 5% NH_4_OH and 50% acetonitrile (ACN) in 5% NH^4^OH, vacuum dried in autosampler vials, and stored at −80 C until LC-MS/MS analysis for phosphoproteomic analysis.

### LC-MS/MS analysis

Unenriched peptides and enriched phosphopeptides from each fraction were analyzed by LC-MS/MS on an Orbitrap Exploris 480 (ThermoFisher, Bremen, Germany) connected to a Dionex 3700 RSLC Nano using 100 µm PicoFrit column self-packed with 2.2 μm Acclaim RSLC 120 C18 (ThermoFisher). A gradient 120-minute LC method was set as: mobile phase A (2% CAN, 0.2% formic acid), mobile phase B (80% ACN, 10% isopropanol, 10% water, 0.2% formic acid), an exponential gradient (curve 6) from 2% to 35% mobile phase B in 100 minutes, followed by a 3 minute linear increase to 45% phase B, a 3 minute linear increase to 90% that was held for two minutes, finishing with a return to 2% phase B that was equilibrated until 120 minutes. Data-dependent acquisition (DDA) was set as: MS1 survey scan data from m/z 340-1800 at 120,000 resolving power (at m/z 120), 300% AGC, max fill time of 100 ms, repeated every 3 seconds; MS/MS scan from m/z 110, a minimum precursor intensity of 70,000, quadrupole isolation width of 0.7 Thompson, 100% AGC target, max fill time of 80 ms, NCE=39, for precursor charge states of 2-4.

### MS data analysis

The mass spectra were searched against a UniProt mouse protein database (version, Jan. 2021) by Andromeda algorithm on the MaxQuant (ver. 1.6.17.0) proteomics analysis platform. The search parameters were set: carbamidomethylation on cysteine residues, TMT-6plex modification on N-terminal and lysine residues as fixed modifications; protein N-terminal acetylation, oxidation on methionine residues and phosphorylation at serine, threonine, and tyrosine residues as variable modifications; a maximum of two missed cleavages. The data were searched against target decoy database and the false discovery rate was set to 1% at the peptide level. Precursor Ion Fraction purity (PIF) was set to 0.75 or higher. The reported ion intensities were used to calculate the changes of proteins and phosphorylation in stimulated CD8+ T cells from *Cd4^Cre^Ppp2ca*^f/f^ mice versus WT mice. Student t-test was used to evaluate significance of changes. A 1.5-fold change with p-value <0.05 was chosen as a cutoff for significantly changed proteins or phosphorylation sites. Ingenuity pathway analysis (IPA, version 107193442) (Qiagen) was used to identify pathways affected by *Ppp2ca* deletion. The volcano plots for the significantly changed proteins or phosphorylation sites were generated using TIBCO Spotfire software, version 12.0.8.

### Metabolic Assay

T cell oxygen consumption rate (OCR) was measured using a Seahorse XFe96 Extracellular Flux Analyzed following established protocols (Agilent). Briefly, equal number of live T cells were seeded at 200,000 cells/well on Cell-Tak (Corning) coated XFe96 plate with fresh XF media (Seahorse XF RPMI medium containing 10 mM glucose, 2 mM L-glutamine, and 1 mM sodium pyruvate, PH 7.4; all reagents from Agilent). OCR was measured in the presence of Oligomycin (1.5 μM; Sigma-Aldrich), FCCP (1.5 μM; Sigma-Aldrich), and Rotenone (1 μM; Sigma-Aldrich)/ Antimycin A (1 μM; Sigma-Aldrich) in Mitostress assay.

## Acknowledgement

We acknowledge the assistance of the Mayo Clinic Proteomics Core, which is a shared resource of the Mayo Clinic Cancer Center (NCI P30 CA15083). The study is supported by NIH grants R01 AI 162678 and R01 AR077518 (for H.Z.).

## Author contributions

X.Z. and H.Z. conceived the project, designed the research, interpreted the data and wrote the manuscript. X.Z., M.L. and M.A. prepare the materials and carried out the experiments. Y.L. managed the mouse colony and performed molecular biology experiments. M.J.H and L.R.P. provided some of the animals and reagents. J.Z., K.L.J., R.Z., and A.P. performed proteomics and phosphoproteomics analysis.

